# Covalent Ligand Screening Uncovers a RNF4 E3 Ligase Recruiter for Targeted Protein Degradation Applications

**DOI:** 10.1101/439125

**Authors:** Carl C. Ward, Jordan I. Kleinman, Scott M. Brittain, Patrick S. Lee, Clive Yik Sham Chung, Kenneth Kim, Yana Petri, Jason R. Thomas, John A. Tallarico, Jeffrey M. McKenna, Markus Schirle, Daniel K. Nomura

**Affiliations:** Department of Molecular and Cell Biology, University of California, Berkeley, Berkeley, CA 94720; Novartis-Berkeley Center for Proteomics and Chemistry Technologies; Department of Chemistry, University of California, Berkeley, Berkeley, CA 94720; Novartis Institutes for BioMedical Research, Cambridge, MA 02139; Novartis Institutes for BioMedical Research, Emeryville, CA 94608; Vertex Pharmaceuticals, Boston, MA 02210; Department of Nutritional Sciences and Toxicology, University of California, Berkeley, Berkeley, CA 94720

**Keywords:** chemoproteomics, RNF4, targeted protein degradation, degraders, covalent ligand, activity-based protein profiling

## Abstract

Targeted protein degradation has arisen as a powerful strategy for drug discovery allowing the targeting of undruggable proteins for proteasomal degradation. This approach most often employs heterobifunctional degraders consisting of a protein-targeting ligand linked to an E3 ligase recruiter to ubiquitinate and mark proteins of interest for proteasomal degradation. One challenge with this approach, however, is that only few E3 ligase recruiters currently exist for targeted protein degradation applications, despite the hundreds of known E3 ligases in the human genome. Here, we utilized activity-based protein profiling (ABPP)-based covalent ligand screening approaches to identify cysteine-reactive small-molecules that react with the E3 ubiquitin ligase RNF4 and provide chemical starting points for the design of RNF4-based degraders. The hit covalent ligand from this screen reacted with either of two zinc-coordinating cysteines in the RING domain, C132 and C135, with no effect on RNF4 activity. We further optimized the potency of this hit and incorporated this potential RNF4 recruiter into a bifunctional degrader linked to JQ1, an inhibitor of the BET family of bromodomain proteins. We demonstrate that the resulting compound CCW 28-3 is capable of degrading BRD4 in a proteasome- and RNF4-dependent manner. In this study, we have shown the feasibility of using chemoproteomics-enabled covalent ligand screening platforms to expand the scope of E3 ligase recruiters that can be exploited for targeted protein degradation applications.

## Main text

Targeted protein degradation is a groundbreaking drug discovery approach for tackling the undruggable proteome by exploiting the cellular protein degradation machinery to selectively eliminate target proteins ^1,2^. This technology most often involves the utilization of heterobifunctional degrader molecules consisting of a substrate-targeting ligand linked to an E3 ligase recruiter. These degraders are capable of recruiting E3 ligases to specific protein targets to ubiquitinate and mark targets for degradation in a proteasome-dependent manner. As functional inhibition of the target is not necessary for degrader efficacy, this strategy has the potential to target and degrade any protein in the proteome for which there exists a ligand. However, while there are ~600 different E3 ligases, only a few E3 ligases that have been successfully exploited in such a strategy, including small-molecule recruiters for cereblon, VHL, MDM2, and cIAP ^2,3^. Identifying facile strategies for discovering ligands that bind to E3 ligases remains crucial for expanding the set of E3 ligase recruiters that can be utilized for targeted protein degradation of a given target of interest and ultimately expand the applicability of this modality to any protein that can be subjected to proteasomal degradation regardless of subcellular localization or tissue-specific expression.

Activity-based protein profiling (ABPP) has arisen as a powerful platform for ligand discovery against targets of interest, including proteins commonly considered as undruggable ^4–10^. ABPP utilizes reactivity-based chemical probes to map proteome-wide reactive and ligandable hotspots directly in complex biological systems ^11,12^. When used in a competitive manner, covalently-acting small-molecule libraries can be screened for competition against the binding of reactivity-based probes to facilitate covalent ligand discovery against proteins of interest ^4–7,9,13–15^. In order to discover covalent ligands that may react with E3 ubiquitin ligases, we first investigated whether representative commercially available E3 ligases -- MDM2, RNF4, and UBE3A -- could be labeled by the cysteine-reactive tetramethylrhodamine-5-iodoacetamide dihydroiodide (IA-rhodamine) reactivity-based probe *in vitro*. We observed IA-rhodamine labeling of all three E3 ligases in a dose-responsive manner **(Fig. 1A)**. While previous studies have already uncovered MDM2 and UBE3A small-molecule modulators ^16–18^, no chemical tools exist for the E3 ubiquitin ligase RNF4, which recognizes SUMOylated proteins and ubiquitinates these proteins for subsequent proteasomal degradation ^19,20^. We thus focused our efforts on developing a ligand for RNF4 as E3 ligase recruitment module for heterbifunctional degraders.

**Figure 1.**
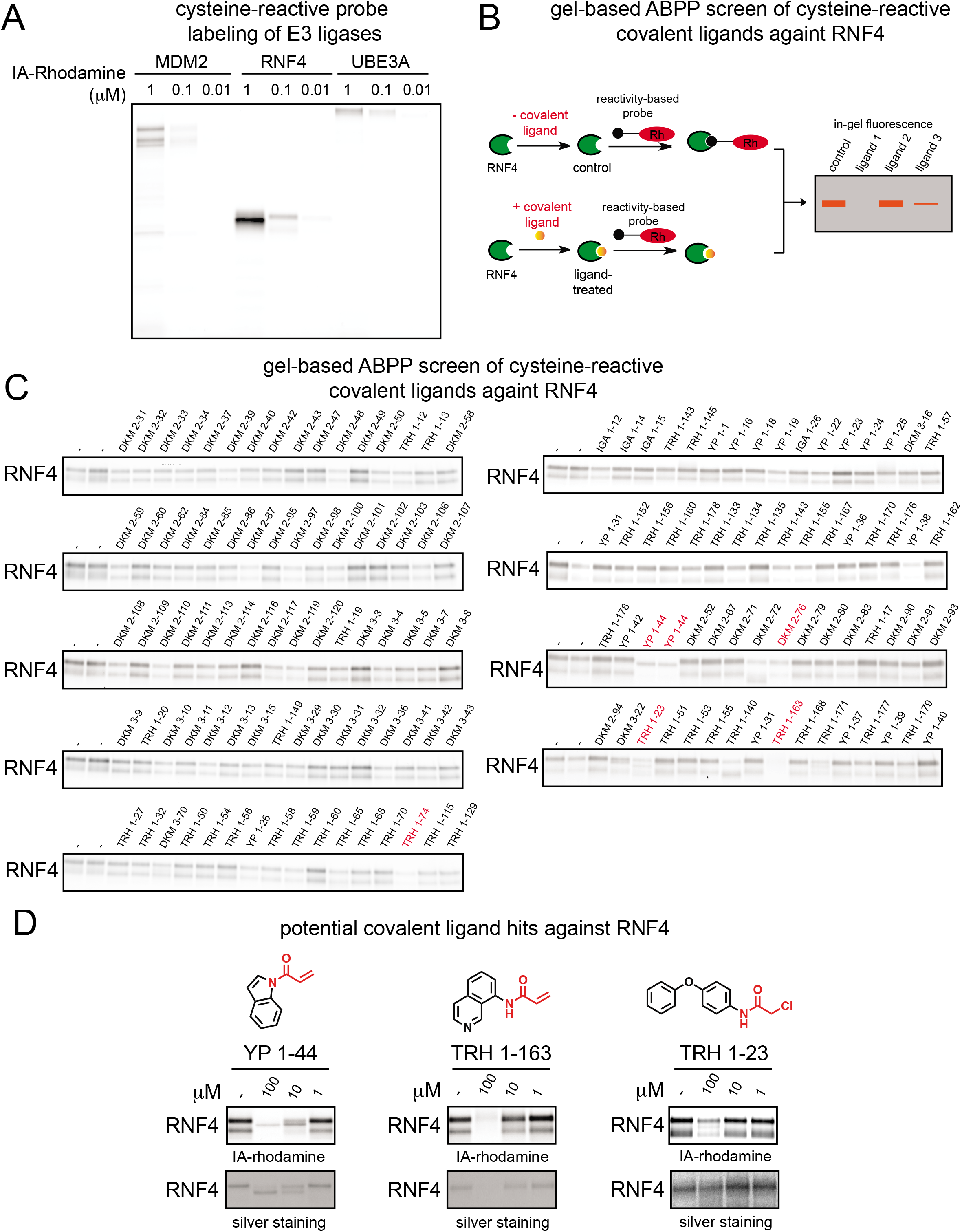
Covalent ligand screen against RNF4 using gel-based ABPP. **(A)** Gel-based ABPP labeling of E3 ligases MDM2, RNF4, and UBE3A. Purified protein was labeled with IA-rhodamine (IA-Rh) for 30 min at room temperature, followed by SDS/PAGE and visualization by in-gel fluorescence. **(B)** Schematic of gel-based ABPP screen of covalent ligands (50 μM) against IA-rhodamine labeling of pure RNF4; **(C)** Gel-based ABPP screen of cysteine-reactive covalent ligands against IA-rhodamine labeling of RNF4. Covalent ligands were pre-incubated with pure RNF4 protein for 30 min prior to IA-rhodamine labeling (250 nM) for 1 h. Proteins were subjected to SDS/PAGE and visualized by in-gel fluorescence. Highlighted in red were potential hits from this screen. **(D)** Chemical structures and gel-based ABPP confirmation of reproducible RNF4 screening hits, performed as described in **(C)**. Gels were also silver stained to identify compounds that induce protein precipitation.

In search of RNF4-targeting ligands, we screened a cysteine-reactive covalent ligand library against IA-rhodamine labeling of pure human RNF4 using gel-based ABPP **(Fig. 1B; Table S1)**. We identified several potential hits from this screen, including TRH 1-74, YP 1-44, DKM 2-76, TRH 1-23, and TRH 1-163. From these, YP 1-44, TRH 1-163, and TRH 1-23 showed reproducible and dose-responsive inhibition of IA-rhodamine labeling of RNF4 **(Fig. 1D, Fig. S1)**. Based on corresponding silver staining of RNF4 in these experiments, we found that TRH 1-163 may be causing protein precipitation. Based on gel-based ABPP analysis of general cysteine-reactivity in 231MFP lysates, YP 1-44 was much less selective compared to TRH 1-23 **(Fig. S1)**. Thus, TRH 1-23 appeared to be the most promising RNF4 hit **(Fig. 1D)**.

We next sought to identify the site-of-modification of TRH 1-23 within RNF4. We performed liquid chromatography-tandem mass spectrometry (LC-MS/MS) analysis of tryptic digests from TRH 1-23-treated purified RNF4 protein and found that TRH 1-23 covalently modified either, but not both, of the two zinc-coordinating cysteines C132 and C135 in the RING domain of RNF4 **(Fig. 2A, 2B)**. In support of its utility as a functional RNF4 recruiter, we wanted to see if TRH 1-23 had any effect upon RNF4 autoubiquitination activity, since previous studies had shown that mutation of both cysteines to serines inhibited RNF4 function ^21–23^. Surprisingly, TRH 1-23 treatment did not inhibit RNF4 autoubiquitination activity in an *in vitro* reconstituted assay **(Fig. 2C)**.

**Figure 2.**
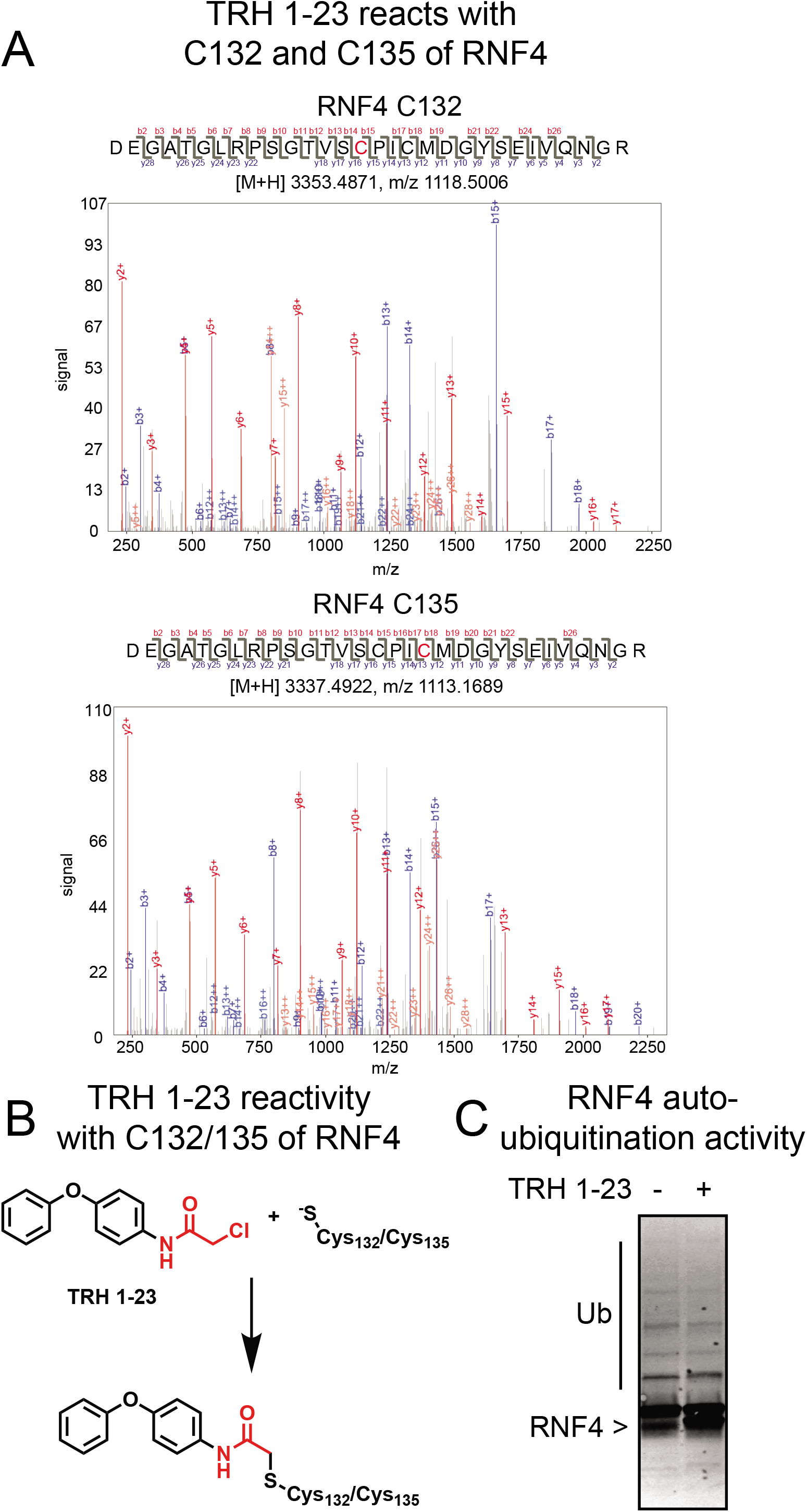
TRH 1-23 reacts non-functionally with zinc-coordinating cysteines in RNF4. **(A)** LC-MS/MS analysis of TRH 1-23 covalent adduct on RNF4. RNF4 was incubated with TRH 1-23 (50 μM) for 30 min at RT. Tryptic digests of RNF4 were analyzed by LC-MS/MS searched for presence of peptides showing the TRH 1-23 adduct. Shown are the MS/MS spectra of TRH 1-23-modified C132 and C135 RNF4 tryptic peptide. Highlighted in red in the peptide sequence is the cysteine that was modified. **(B)** Schematic of TRH 1-23 reactivity with C132 or C135 of RNF4. **(C)** TRH 1-23 does not inhibit RNF4 autoubiquitination. RNF4 was preincubated with TRH 1-23 (100 μM) for 30 min at room temperature followed by addition of UBA1, E2 enzyme, and ATP for 40 min at 37 C. The reaction was quenched and subjected to SDS/PAGE and Western blotting for RNF4. Gel shown in **(C)** is a representative gel from n=3.

While a promising non-functional ligand against RNF4, we considered the potency of TRH 1-23 with a 50% inhibitory concentration (IC50) in the double digit μM range in competitive ABPP suboptimal for usage as an efficient RNF4 recruiter. We thus synthesized several TRH 1-23 analogs and used gel-based ABPP to test their potency and structure-activity relationships against RNF4 **(Fig. 3A)**. Among these analogs, we found CCW 16, with an *N*-benzyl and 4-methoxyphenoxyphenyl substitution on the chloroacetamide scaffold, to be among the most potent of the analogs with inhibition of IA-rhodamine labeling of RNF4 observed down to 1 μM. Further confirmatory studies revealed the IC50 for CCW 16 to be 1.8 μM **(Fig. 3B)**.

**Figure 3.**
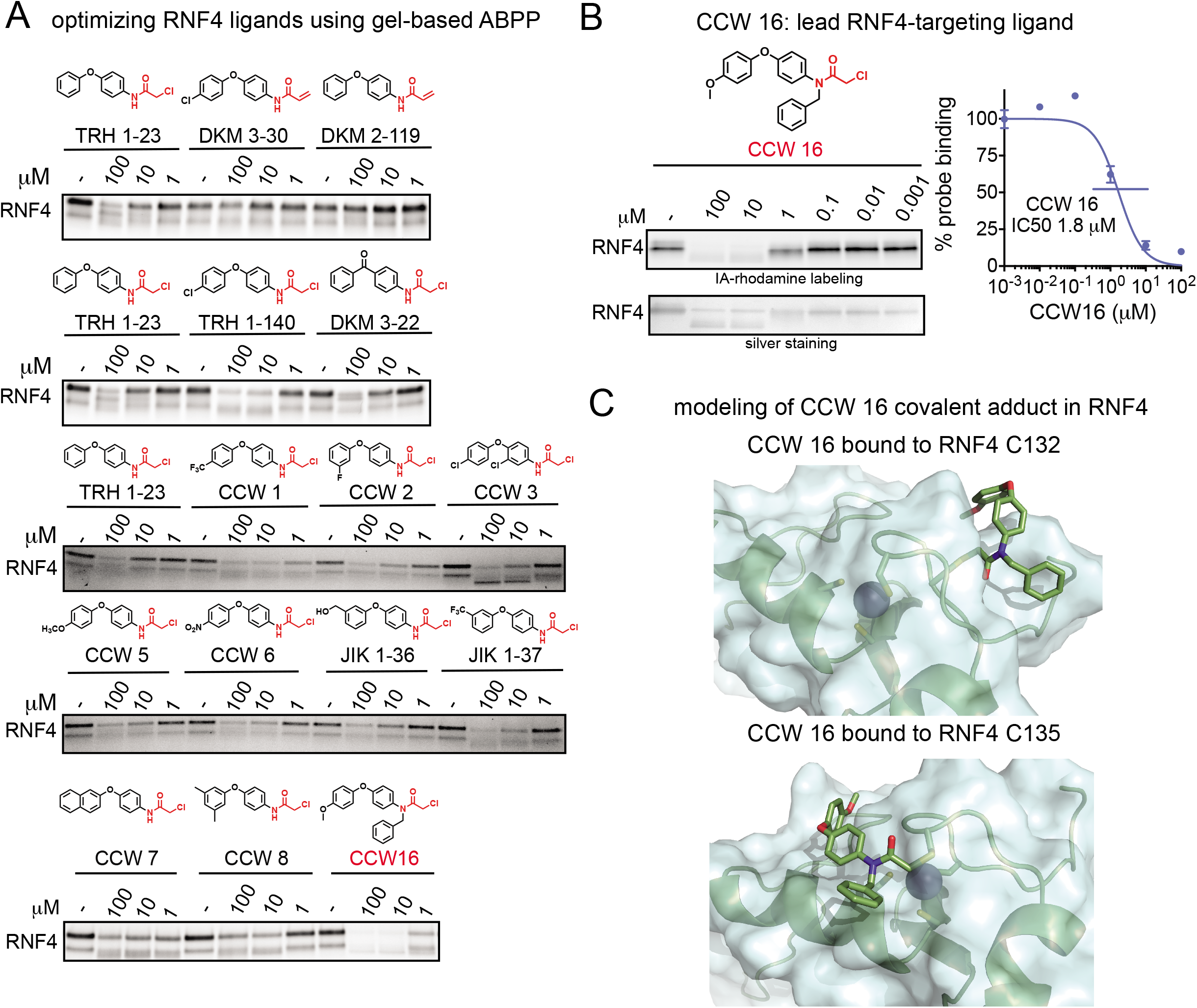
Optimizing RNF4 covalent ligands using gel-based ABPP. **(A)** Derivatives of TRH 1-23 were tested against IA-rhodamine labeling of RNF4 using gel-based ABPP. Silver stained gels are also shown to visualize total protein content. **(B)** CCW16 was tested against IA-rhodamine labeling of RNF4 using gel-based ABPP. For **(A)** and **(B)**, covalent ligands were pre-incubated with pure RNF4 protein for 30 min prior to IA-rhodamine labeling for 1 h. Proteins were subjected to SDS/PAGE and visualized by in-gel fluorescence. In **(B)**, gels were quantified by densitometry to calculate IC50 values. Gel shown in **(B)** is a representative gel from n=3. **(C)** Covalent docking of CCW 16 bound to either C132 or C135 in RNF4.

To better understand how CCW 16 was interacting with RNF4, we carried out covalent docking on CCW 16 bound to either C132 or C135 on a publicly available crystal structure of RNF4 (PDB: 4PPE) **(Fig. 3C)**. In this model, CCW 16 covalent binding to C132 caused a small portion of the backbone to rotate flipping the C132 sidechain out into a more surface accessible conformation. The CCW 16 ligand occupied a groove on the RNF4 surface consisting of side-chains R125, T129, and S131, with no obvious polar interactions between the ligands and the protein in this model. Interestingly, CCW 16 did not occupy the zinc-binding site, left the other three zinc-coordinating cysteines (C135, C159, and C162) unperturbed (**Fig. 3C**). In modeling CCW 16 covalently bound to C135, we observed a similar association with a surface groove of RNF4 occupying a region between Q161, S166, and P174. However, to accommodate the ligand, only a dihedral rotation of Cα & Cβ of C132 was predicted as necessary **(Fig. 3C)**. Both models indicate that zinc may still be bound by the three remaining cysteines in the site, while CCW 16 binds outside of the zinc-coordinating site. Since CCW 16 and TRH 1-23 are based on the same scaffold, these results may explain why modification by TRH 1-23 did not inhibit RNF4 autoubiquitination activity while the reported mutation of both cysteines 132 and 135 to serine leads to loss of activity.

Based on the structure-activity relationships observed with TRH 1-23 analogs, the 4-methoxy group on CCW16 presented an ideal position for extending a linker to yield a RNF4-based degrader. To demonstrate that we could use CCW 16 as a potential RNF4 recruiter for targeted protein degradation applications, we synthesized CCW 28-3, a bifunctional degrader linking CCW 16 to the BET bromodomain family inhibitor JQ1 **(Fig. 4A)** ^24^. CCW 28-3 was prepared in five steps from commercially available materials. Demethylation of 4-(4-methoxyphenoxy)aniline yielded 4-(4-aminophenoxy)phenol, which underwent reductive amination with benzaldehyde using sodium triacetoxyborohydride as the reducing agent to form CCW 22. Alkylation of phenolic moiety of CCW 22 with 4-(Boc-amino)butyl bromide and subsequent reaction with 2-choloracetyl chloride allowed functionalization of the RNF4 recruiter with a linker containing a “latent” reactive handle, a Boc-protected amino group. Boc deprotection by trifluoroacetic acid restored the functional primary amine which could then be reacted with a hydrolyzed free-acid form of JQ1 through amide coupling, yielding the bifunctional degrader CCW 28-3 as the final product **(Supporting Methods)**. It is noteworthy that the amide coupling reaction should be highly versatile and thus this synthetic scheme should be applicable for conjugating different protein-targeting ligands with carboxylic acid moiety onto our RNF4 recruiter.

**Figure 4.**
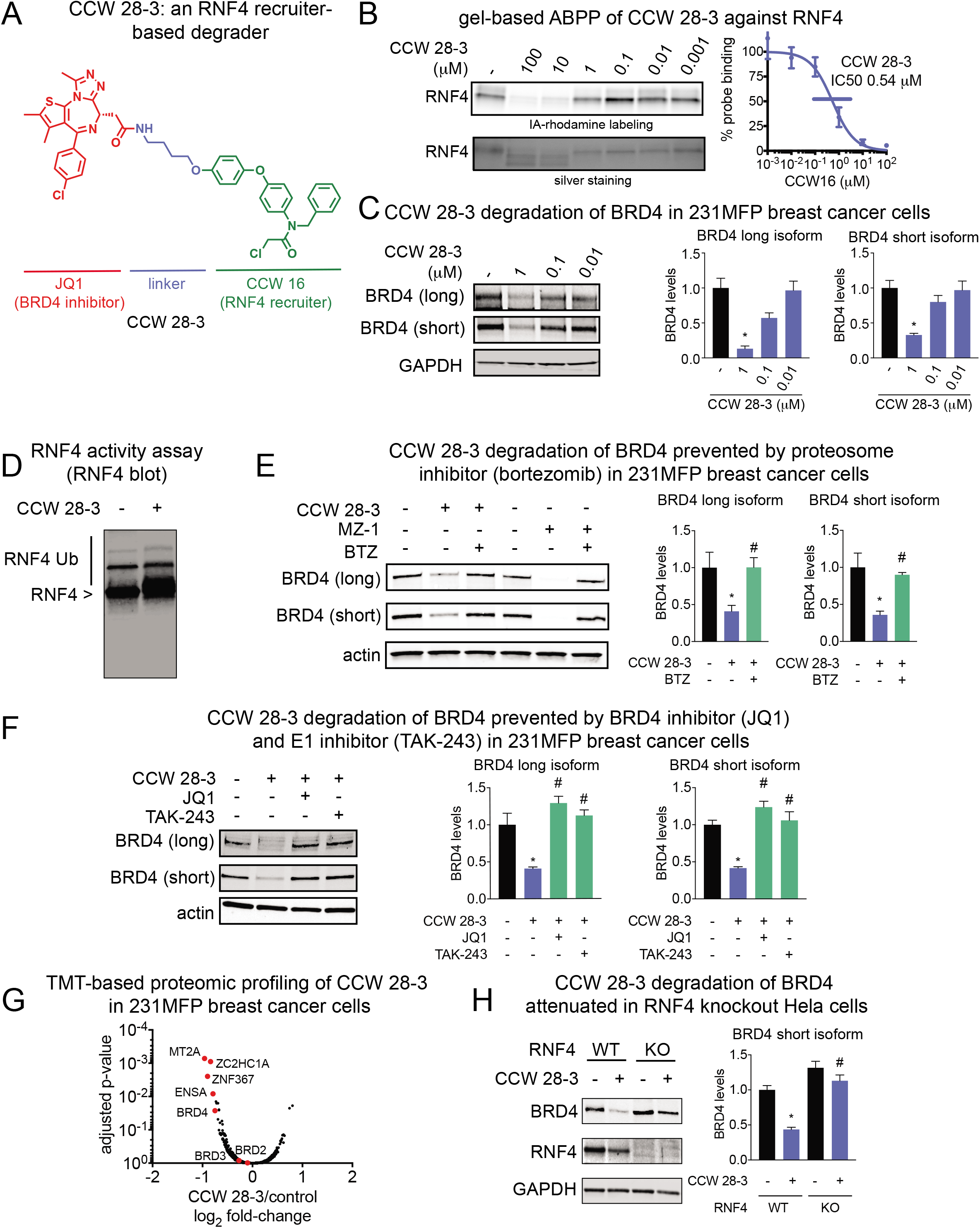
RNF4 recruiter-based BRD4 degrader. **(A)** Structure of CCW 28-3, an RNF4-recruiter-based degrader linked to BRD4 inhibitor JQ1. **(B)** Gel-based ABPP analysis of CCW 28-3 against pure human RNF4. CCW 28-3 was pre-incubated with pure RNF4 protein for 30 min prior to IA-rhodamine labeling for 1 h. Proteins were subjected to SDS/PAGE and visualized by in-gel fluorescence. Gels were quantified by densitometry to calculate IC50 values. **(C)** CCW 28-3 treatment in 231MFP breast cancer cells leads to BRD4 degradation. 231MFP breast cancer cells were treated with vehicle DMSO or CCW 28-3 for 3 h. Proteins were subjected to SDS/PAGE and Western blotting for BRD4 and GAPDH loading control. **(D)** CCW 28-3 does not inhibit RNF4 autoubiquitination. RNF4 was pre-incubated with CCW 28-3 (10 μM) for 30 min at room temperature followed by addition of UBA1, E2 enzyme, and ATP for 60 min at 37 C. The reaction was quenched and subjected to SDS/PAGE and Western blotting for RNF4. **(E, F)** CCW 28-3 treatment in 231MFP breast cancer cells leads to proteasome-, E1 inhibitor-, and BRD4 inhibitor-dependent BRD4 degradation. Vehicle DMSO or proteasome inhibitor bortezomib (BTZ) (10 μM) **(E)**, E1 inhibitor TAK-243 (10 μM), or BRD4 inhibitor JQ1 (10 μM) **(F)** were pre-incubated for 30 min prior to treatment with MZ1 (1 μM) or CCW 28-3 (1 μM) for 3 h. Proteins were subjected to SDS/PAGE and Western blotting for BRD4 and actin loading control. **(G)** TMT-based quantitative proteomic analysis of protein expression changes from CCW 28-3 (1 μM) treatment for 3 h in 231MFP cells. **(H)** RNF4 wild-type and knockout Hela cells were treated with CCW 28-3 (10 μM) for 5 h and subjected to SDS/PAGE and Western blotting for BRD4, RNF4, and GAPDH. Blots in **(B-F, H)** were quantified by densitometry. Data in **(B-F, H)** are from representative gels from n=3. Bar graphs are average ± sem, n=3-5/group. Significance is expressed as *p<0.05 compared to vehicle-treated controls and #p<0.05 compared to CCW 28-3 treated groups in **(E-F)** and CCW 28-3 treated wild-type cells in **(H)**. Data in (**G**) represent 6180 protein groups quantified with 2 or more unique peptides in triplicate treatments, see **Table S2** for details.

CCW 28-3 showed higher potency for RNF4 than CCW 16 with an IC50 of 0.54 μM in the competitive ABPP assay **(Fig. 4B)**. Compellingly, treatment with CCW 28-3 degraded BRD4 in a dose-responsive manner in 231MFP breast cancer cells **(Fig. 4C)**. We also confirmed that CCW 28-3 did not inhibit RNF4 autoubiquitination activity **(Fig. 4D)**. CCW 28-3-mediated degradation of BRD4 was prevented by pretreatment of cells with the proteasome inhibitor bortezomib (BTZ), JQ1 alone, as well as the E1 ubiquitin activating enzyme inhibitor TAK-243 **(Fig. 4E-4F)**. We next performed tandem mass tagging (TMT)-based quantitative proteomic profiling to determine the selectivity of changes in protein expression from CCW 28-3 treatment. We showed that BRD4 is one of the primary targets degraded by CCW 28-3 in 231MFP breast cancer cells **(Fig. 4G; Table S2)**. We also observed additional downregulated targets such as MT2A, ZC2HC1A, ZNF367, and ENSA, which may represent potential off-targets of JQ1 or transcriptional or post-translationally-driven changes in protein expression stemming from on or off-target effects of our RNF4-targeting ligand in 231MFP cells. Interestingly, CCW 28-3 treatment did not lead to BRD2 and 3 degradation. Previously reported JQ1-based degraders have shown varying levels of degradation of BET bromodomain family members with Cereblon and VHL-recruiting modules ^25–28^. This study highlights the utility of having additional E3 ligase recruiters for tuning the efficacy and selectivity of degraders targeting a given protein of interest.

We next used isotopic tandem orthogonal proteolysis-enabled ABPP (isoTOP-ABPP) platforms to assess the proteome-wide selectivity of CCW 28-3 ^4,6,7,29^ **(Fig. S2; Table S2)**. We treated 231MFP cells with vehicle or CCW 28-3 and labeled the resulting proteomes with the cysteine-reactive *N*-hex-5-ynyl-2-iodoacetamide (IA-alkyne) probe, followed by appendage of isotopically light or heavy TEV protease-cleavable biotin-azide tags onto probe-labeled proteins in vehicle and CCW 28-3-treated groups, respectively. Probe-modified peptides were enriched, eluted, and analyzed using previously described methods for isoTOP-ABPP ^4,6,7,29^. Through this analysis, we demonstrated that out of 1114 total quantified probe-modified peptides only 7 potential off-targets of CCW 28-3 showed isotopically light to heavy ratios greater than 4. This ratio >4 indicated that the covalent ligand displaced IA-alkyne probe labeling at the particular site within the protein by >75 %. Most notably, none of these off-targets of CCW 28-3 are part of the ubiquitin-proteasome system indicating that the observed BRD4 degradation can likely be attributed to RNF4-based ubiquitination **(Fig. S2; Table S2)**. Unfortunately, we were not able to observe the probe-modified peptide for RNF4 in this experiment likely due to its low abundance or labeling efficiency compared to other IA-alkyne labeled proteins. We were thus not able to confirm the degree of RNF4 occupancy by CCW 28-3 using isoTOP-ABPP.

We thus aimed to confirm engagement of RNF4 by CCW 28-3 *in situ* by other means. Since we had shown that CCW 16 was an optimized covalent ligand against RNF4, we synthesized an alkyne-functionalized derivative of CCW 16—CCW 36 **(Fig. S3)**. We then overexpressed RNF4 in HEK293T cells, treated these cells with vehicle or CCW 28-3, enriched for RNF4, and then labeled the enriched RNF4 with CCW 36, followed by CuAAC-mediated appendage of rhodamine-azide to visualize CCW 36 cysteine-reactivity of RNF4 by in-gel fluorescence. Intriguingly, we only observed ގ30 % engagement of RNF4 by CCW 28-3 *in situ* at a concentration 10-times higher than the concentration where we observed significant BRD4 degradation **(Fig. S3; Fig 4C, E, F)**. We attribute the low degree of target engagement to poor cell permeability of our yet-unoptimized RNF4-targeting ligand. Nonetheless, our results encouragingly suggest that only a modest degree of engagement of RNF4 is sufficient to facilitate the degradation of BRD4 with CCW 28-3.

Because CCW 28-3 was not completely selective and we were targeting conserved zinc-coordinating cysteines across the RING family of E3 ligases, we next sought to confirm the contributions of RNF4 to CCW 28-3-mediated degradation of BRD4. We compared CCW 28-3-mediated BRD4 degradation in wild-type (WT) and RNF4 knockout (KO) HeLa cells. Convincingly, CCW 28-3-mediated degradation of BRD4 observed in HeLa WT cells was not evident in RNF4 KO cells **(Fig. 4H; Fig. S4)**. These data further support our claim that CCW 28-3 degrades BRD4 specifically through RNF4 recruitment.

In conclusion, our study demonstrates the feasibility of using ABPP-based covalent ligand screening in vitro to rapidly discover chemical entry points for targeting E3 ligases and that these covalent ligand hits can be identified, optimized, and incorporated into degraders for targeted protein degradation applications. While CCW 16 and CCW 28-3 are not yet completely selective for RNF4 and can only achieve fractional target engagement in cells, we demonstrate that we can still degrade BRD4 in a RNF4-dependent manner. We note that CCW 28-3 does not degrade BRD4 as well as other previously reported BRD4 degraders such as MZ1 that utilizes a VHL-recruiter linked to JQ1^25^. Future medicinal chemistry efforts can be employed to optimize the potency, selectivity, and cell permeability of CCW 16 for targeting RNF4 and to improve linker positioning and composition of CCW 28-3 to promote better degradation of protein substrates. Nonetheless, CCW 16 represents a novel, small-molecule E3 ligase recruiter for RNF4, beyond the four other E3 ligase recruiters that have been reported previously, targeting cereblon, VHL, MDM2, and cIAP ^2^. Beyond RNF4, our study also highlights zinc-coordinating cysteines as a potential ligandable modality that can be targeted with cysteine-targeting covalent ligands. We believe that the approaches described here, and insights gained in this study can be utilized for future applications in expanding the scope of E3 ligase recruiters or modulators.

## Methods

### Covalent Ligand Library used in Initial Screen

The synthesis and characterization of many of the covalent ligands screened against RNF4 have been previously reported ^4–6,13^. Synthesis of TRH 1-156, TRH 1-160, TRH 1-167, YP 1-16, YP 1-22, YP 1-26, YP 1-31, YP 1-44 have been previously reported ^30–37^. The synthesis and characterization of covalent ligands that have not been reported are described in **Supporting Information**.

### Gel-Based ABPP

Gel-based ABPP methods were performed as previously described ^5,6,38,39^. Pure recombinant human RNF4 was purchased from Boston Biochem (K-220). RNF4 (0.25 μg) was diluted into 50 μL of PBS and 1μL of either DMSO (vehicle) or covalently acting small molecule to achieve the desired concentration. After 30 minutes at room temperature, the samples were treated with 250 nM IA-Rhodamine (Setareh Biotech, 6222, prepared in anhydrous DMSO) for 1 h at room temperature. Samples were then diluted with 20 μL of 4x reducing Laemmli SDS sample loading buffer (Alfa Aesar) and heated at 90 °C for 5 min. The samples were separated on precast 4-20% Criterion TGX gels (Bio-Rad Laboratories, Inc.). Fluorescent imaging was performed on a ChemiDoc MP (Bio-Rad Laboratories, Inc) inhibition of target labeling was assessed by densitometry using ImageJ.

### LC-MS/MS analysis of RNF4

Purified RNF4 (10 μg) was diluted into 80 μL of PBS and treated for 30 min with DMSO or compound (50 μM). The DMSO control was then treated with light iodoacetamide (IA) while the compound treated sample was incubated with heavy IA for 1 h each at room temperature (100 μM, Sigma-Aldrich, 721328). The samples were precipitated by additional of 20 μL of 100% (w/v) TCA and combined pairwise before cooling to −80 C for one hour. The combined sample was then spun for at max speed for 20 min at 4 °C, supernatant is carefully removed and the sample is washed with ice cold 0.01 M HCl/90 % acetone solution. The sample was then resuspended in 2.4 M urea containing 0.1 % Protease Max (Promega Corp. V2071) in 100 mM ammonium bicarbonate buffer. The samples were reduced with 10 mM TCEP at 60 °C for 30 min. The samples were then diluted 50% with PBS before sequencing grade trypsin (1 ug per sample, Promega Corp, V5111) was added for an overnight incubation at 37 °C. The next day the sample was centrifuged at 13200 rpm for 30 min. The supernatant was transferred to a new tube and acidified to a final concentration of 5 % formic acid and stored at −80 °C until MS analysis.

### RNF4 ubiquitination assay

For *in vitro* auto-ubiquitination assay, 200 nM RNF4 in 15 μL ubiquitination assay buffer (50 mM Tris, 150mM NaCl, 5 mM MgCl_2_, 5mM DTT, pH 7.4) was pre-incubated with DMSO vehicle or the covalently-acting compound for 30 min at room temperature. Subsequently, UBE1 (50 nM, Boston Biochem, E-305), UBE2D1 (400nM Boston Bichem, E2-615), Flag-ubiquitin (4000 nM, Boston Biochem, U-120) and ATP (200 μM) were added in ubiquitination assay buffer bring the total volume to 30 μL. The mixture was incubated at RT for 30 min before quenching with 10 μL of 4x Laemmli’s buffer. Ubiquitination activity was measured by separation on an SDS-PAGE gel and western blotting as previously described.

### Synthetic Methods and Characterization of Covalent Ligand Analogs and CCW 28-3 Degrader

Synthetic methods and characterization are detailed in Supporting Information

### Covalent Docking of CCW 16 in RNF4

For covalent docking, a crystal structure of human RNF4 (PDB code: 4PPE) was used ^40^. This crystal structure was then prepared for docking utilizing Schrödinger’s Maestro (2018-1) protein preparation ^41^. Missing loops and side chains were added using PRIME and only A chain was utilized for docking purposes. Protonation was carried out to optimize H-bond assignments (assuming pH 7.0) and all waters were removed. A restrained minimization (<0.3Å) was then carried out to optimize the protein. A Zn coordinated by C132, C135, C159 & C162 was removed for docking purposes.

Prior to docking, CCW16 was prepared via LigPrep. To carry out covalent docking using Schrödinger’s covalent docking ^42^, either C132 or C135 were defined as the center of the binding grid (within 20A). CCW 16 was selected as the ligand and the appropriate reactive residues were selected. A “nucleophilic substitution” was selected as the reaction type and calculations were carried out on a Linux workstation with Intel Xeon 2.4GHz processors running Red Hat 6.8 with 128GB memory.

### Cell Culture

The 231MFP cells were obtained from Prof. Benjamin Cravatt and were generated from explanted tumor xenografts of MDA-MB-231 cells as previously described^43^. RNF4 knockout HeLa cells were purchased from EdiGene USA (CL0033025003A). RNF4 wild-type HeLa cells were provided by EdiGene USA or the UC Berkeley Cell Culture Facility. 231MFP cells were cultured in L-15 media (Corning) containing 10% (v/v) fetal bovine serum (FBS) and maintained at 37°C with 0% CO_2_. HeLa cells were cultured in DMEM media (Corning) containing 10% (v/v) fetal bovine serum (FBS) and maintained at 37°C with 5% CO_2_.

### Cell based degrader assays

For assaying degrader activity, cells were seeded (500,000 for 231MFP cells, 300,000 for HeLa cells) into a 6 cm tissue culture dish (Corning) in 2.0 – 2.5 mL of media and allowed to adhere overnight. The following morning, media was replaced with complete media containing the desired concentration of compound diluted from a 1000x stock in DMSO. At the specified timepoint, cells were washed once with PBS on ice, before 150uL of lysis buffer was added to the plate (10 mM sodium phosphate, 150 mM NaCl, 0.1% SDS, 0.5% sodium deoxycholate, 1% Triton X100). The cells were incubated in lysis buffer for 5 min before scraping and transfer to microcentrifuge tubes. The lysates were then frozen at −80C or immediately processed for western blotting. To prepare for western blotting, the lysates were cleared with 20,000 g spin for 10 min and the resulting supernatants quantified via BCA assay. The lysates were normalized by dilution with PBS to match the lowest concentration lysate and appropriate amount of 4x Laemmli’s reducing buffer added.

### Western blotting

Antibodies to RNF4 (Proteintech, 17810-1-AP, 1:1000), GAPDH (Proteintech, 60004-1-IG, 1:5000), BRD4 (Abcam, Ab128874, 1:1000), and beta-actin (Proteintech Group Inc., 6609-1-IG, 1:7000) were obtained from the specified commercial sources and dilutions were prepared in 5% BSA/TBST at the specified dilutions. Proteins were resolved by SDS/PAGE and transferred to nitrocellulose membranes using the iBlot system (Invitrogen). Blots were blocked with 5 % BSA in Tris-buffered saline containing Tween 20 (TBST) solution for 1 h at room temperature, washed in TBST, and probed with primary antibody diluted in recommended diluent per manufacturer overnight at 4 °C. Following washes with TBST, the blots were incubated in the dark with secondary antibodies purchased from Ly-Cor and used at 1:10,000 dilution in 5 % BSA in TBST at room temperature. Blots were visualized using an Odyssey Li-Cor scanner after additional washes. If additional primary antibody incubations were required the membrane was stripped using ReBlot Plus Strong Antibody Stripping Solution (EMD Millipore, 2504), washed and blocked again before being re-incubated with primary antibody.

### IsoTOP-ABPP chemoproteomic studies

IsoTOP-ABPP studies were done as previously reported ^4,6,7,29^. Briefly, cells were lysed by probe sonication in PBS and protein concentrations were measured by BCA assay ^44^. For *in situ* experiments, cells were treated for 90 min with either DMSO vehicle or covalently-acting small molecule (from 1000X DMSO stock) before cell collection and lysis. For *in vitro* experiments, proteome samples diluted in PBS (4 mg of proteome per biological replicate) were treated with a DMSO vehicle or covalently-acting small molecule for 30 min at room temperature. Proteomes were subsequently treated with IA-alkyne (100 μM, Chess GmbH, 3187) for 1 h at RT. CuAAC was performed by sequential addition of tris(2-carboxyethyl)phosphine (TCEP) (1 mM, Sigma), tris[(1-benzyl-1H-1,2,3-triazol-4-yl)methyl]amine (TBTA) (34 μM, Sigma), copper (II) sulfate (1 mM, Sigma), and biotin-linker-azide—the linker functionalized with a TEV protease recognition sequence as well as an isotopically light or heavy valine for treatment of control or treated proteome, respectively. After CuAAC, proteomes were precipitated by centrifugation at 6500 × g, washed in ice-cold methanol, combined in a 1:1 control/treated ratio, washed again, then denatured and resolubilized by heating in 1.2 % SDS/PBS to 80°C for 5 minutes. Insoluble components were precipitated by centrifugation at 6500 × g and soluble proteome was diluted in 5 ml 0.2% SDS/PBS. Labeled proteins were bound to avidin-agarose beads (170 μl resuspended beads/sample, Thermo Pierce) while rotating overnight at 4°C. Bead-linked proteins were enriched by washing three times each in PBS and water, then resuspended in 6 M urea/PBS (Sigma) and reduced in TCEP (1 mM, Sigma), alkylated with iodoacetamide (IA) (18 mM, Sigma), then washed and resuspended in 2 M urea and trypsinized overnight with 2 ug/sample sequencing grade trypsin (Promega). Tryptic peptides were eluted off.

Beads were washed three times each in PBS and water, washed in TEV buffer solution (water, TEV buffer, 100 μM dithiothreitol) and resuspended in buffer with Ac-TEV protease (Invitrogen) and incubated overnight. Peptides were diluted in water and acidified with formic acid (1.2 M, Spectrum) and prepared for analysis.

### IsoTOP-ABPP Mass Spectrometry Analysis

Peptides from all chemoproteomic experiments were pressure-loaded onto a 250 μm inner diameter fused silica capillary tubing packed with 4 cm of Aqua C18 reverse-phase resin (Phenomenex # 04A-4299) which was previously equilibrated on an Agilent 600 series HPLC using gradient from 100% buffer A to 100% buffer B over 10 min, followed by a 5 min wash with 100% buffer B and a 5 min wash with 100% buffer A. The samples were then attached using a MicroTee PEEK 360 μm fitting (Thermo Fisher Scientific #p-888) to a 13 cm laser pulled column packed with 10 cm Aqua C18 reverse-phase resin and 3 cm of strong-cation exchange resin for isoTOP-ABPP studies. Samples were analyzed using an Q Exactive Plus mass spectrometer (Thermo Fisher Scientific) using a 5-step Multidimensional Protein Identification Technology (MudPIT) program, using 0%, 25%, 50%, 80%, and 100% salt bumps of 500 mM aqueous ammonium acetate and using a gradient of 5-55% buffer B in buffer A (buffer A: 95:5 water:acetonitrile, 0.1% formic acid; buffer B 80:20 acetonitrile:water, 0.1% formic acid). Data was collected in data-dependent acquisition mode with dynamic exclusion enabled (60 s). One full MS (MS1) scan (400-1800 m/z) was followed by 15 MS2 scans (ITMS) of the nth most abundant ions. Heated capillary temperature was set to 200 °C and the nanospray voltage was set to 2.75 kV.

Data was extracted in the form of MS1 and MS2 files using Raw Extractor 1.9.9.2 (Scripps Research Institute) and searched against the Uniprot human database using ProLuCID search methodology in IP2 v.3 (Integrated Proteomics Applications, Inc) ^45^. Cysteine residues were searched with a static modification for carboxyaminomethylation (+57.02146) and up to three differential modifications for methionine oxidation and either the light or heavy TEV tags (+464.28596 or +470.29977, respectively). Peptides were required to have at least one tryptic end and to contain the TEV modification. ProLUCID data was filtered through DTASelect to achieve a peptide false-positive rate below 5%. Only those probe-modified peptides that were evident across all two out of three biological replicates were interpreted for their isotopic light to heavy ratios. Those probe-modified peptides that showed ratios >3 were further analyzed as potential targets of the covalently-acting small-molecule. For modified peptides with ratios >3, we filtered these hits for peptides were present in all three biological replicates. For those probe-modified peptide ratios >3, only those peptides with 3 ratios >3 were interpreted, and otherwise replaced with the lowest ratio. For those probe-modified peptide ratios >4, only those peptides with 3 ratios >4 were interpreted, and otherwise replaced with the lowest ratio. MS1 peak shapes of any resulting probe-modified peptides with ratios >3 were then manually confirmed to be of good quality for interpreted peptides across all biological replicates.

### Vectors for RNF4 Overexpression

The pCMV6-Entry-RNF4 (C-term FLAG + Myc tag) vector was purchased from Origene (RC207273). The corresponding pCMV6-Entry-eGFP vector was constructed via Gibson Assembly with primers: GATCTGCCGCCGCGATCGCCatggtgagcaagggcgag, TCGAGCGGCCGCGTACGCGTcttgtacagctcgtccatgcc to amplify the eGFP ORF with desired overlaps, and ACGCGTACGCGGCCG, GGCGATCGCGGCGG to linearize the pCMV6-Entry backbone.

### RNF4 Overexpression, Immunoprecipitation, and *in vitro* Probe Labeling Experiments

HEK293T cells were seeded to 30-40% confluency in 10 cm dishes. The day of transfection, media was replaced with DMEM containing 2.5% FBS. For transfection, 500 μL of Opti-MEM (Thermo Fisher) containing 10 μg of pCMV6-Entry-RNF4 or eGFP vector and 30 ug of polyethylenimine was added to the plate. Media was replaced 48 h later with identical media containing DMSO or CCW 28-3 (10 μM). After 1.5 h, cells were washed and harvested in PBS, resuspended in 500 μL PBS, and lysed by probe tip sonication at 15 % amplitude for 2 × 10 s. Lysates were cleared by centrifugation at 21,000 g for 20 min, and resulting supernatant was mixed with 30 μL of anti-FLAG resin (Genscript, L00432) and rotated at 4 ^°^C for 1 h. The beads were washed 3 × with 500 μL PBS containing 300 mM sodium chloride, before elution of FLAG-tagged proteins with 100 μL of 250 ng/μL 3 × FLAG peptide (APExBIO, A6001) in PBS.

For labeling of enriched RNF4 with CCW 36 to monitor *in situ* CCW 28-3 target engagement, 25 μL of eluted protein was treated with 1 μM CCW 36 (alkyne-functionalized probe of CCW 16) for 1 h. A rhodamine ligand was appended with copper catalyzed click chemistry by addition of 1 mM Copper (II) Sulfate, 35 μM Tris(benzyltriazole methylamine) (TBTA), 1 mM TCEP, and 0.1 mM Azide-Fluor 545 (Click Chemistry Tools, AZ109-5). Degree of labeling was assessed SDS/PAGE and measuring in-gel fluorescence using Image J.

### TMT-based Quantitative Proteomic Analysis

#### Cell Lysis, Proteolysis and Isobaric Labeling

Treated cell-pellets were lysed and digested using the commercially available Pierce™ Mass Spec Sample Prep Kit for Cultured Cells (Thermo Fisher Scientific, P/N 84840) following manufacturer’s instructions. Briefly, 100 μg protein from each sample was reduced, alkylated, and digested overnight using a combination of Endoproteinase Lys-C and trypsin proteases. Individual samples were then labeled with isobaric tags using commercially available Tandem Mass Tag™ 6-plex (TMTsixplex™) (Thermo Fisher Scientific, P/N 90061) or TMT11plex (TMT11plex™) isobaric labeling reagent (Thermo Fisher Scientific, P/N 23275) kits, in accordance with manufacturer’s protocols.

#### High pH Reversed Phase Separation

Tandem mass tag labeled (TMT) samples were then consolidated, and separated using high-pH reversed phase chromatography (RP-10) with fraction collection as previously described ^46^. Fractions were speed-vac dried, then reconstituted to produce 24 fractions for subsequent on-line nanoLC-MS/MS analysis.

#### Protein Identification and Quantitation by nanoLC-MS/MS

Reconstituted RP-10 fractions were analyzed on a Thermo Orbitrap Fusion Lumos Mass Spectrometer (Xcalibur 4.1, Tune Application 3.0.2041) coupled to an EasyLC 1200 HPLC system (Thermo Fisher Scientific). The EasyLC 1200 was equipped with a 20 μL loop, setup for 96 well plates. A Kasil-fritted trapping column (75 μm ID) packed with ReproSil-Pur 120 C18-AQ, 5 μm material (15mm bed length) was utilized together with a 160mm length, 75 μm inner diameter spraying capillary pulled to a tip diameter of approximately 8-10 μm using a P-2000 capillary puller (Sutter Instruments, Novato, CA). The 160mm separation column was packed with ReproSil-Pur 120 C18-AQ, 3 μm material (Dr. Maisch GmbH, Ammerbuch-Entringen, Germany). Mobile phase consisted of A= 0.1% formic acid/2% acetonitrile (v/v), and Mobile phase B= 0.1% formic acid/98% acetonitrile (v/v). Samples (18 μL) were injected on to trapping column using Mobile Phase A at a flow rate of 2.5 μL/min. Peptides were then eluted using an 80 minute gradient (2% Mobile Phase B for 5 min, 2%-40% B from 5-65 min, followed by 70% B from 65-70 minutes, then returning to 2% B from 70-80 min) at a flowrate of 300 nL/min on the capillary separation column with direct spraying into the mass spectrometer. Data was acquired on Orbitrap Fusion Lumos Mass Spectrometer in data-dependent mode using synchronous precursor scanning MS^3^ mode (SPS-MS3), with MS^2^ triggered for the 12 most intense precursor ions within a mass-to-charge ratio *(m/z*) range of 300-1500 found in the full MS survey scan event. MS scans were acquired at 60,000 mass resolution (*R*) at *m/z* 400, using a target value of 4 × 10^5^ ions, and a maximum fill time of 50 ms. MS^2^ scans were acquired as CID ion trap (IT) rapid type scans using a target value of 1 × 10^4^ ions, maximum fill time of 50 ms, and an isolation window of 2 Da. Data-dependent MS^3^ spectra were acquired as Orbitrap (OT) scans, using Top 10 MS^2^ daughter selection, automatic gain control (AGC) target of 5 × 10^4^ ions, with scan range of *m/z* 100-500. The MS^3^ maximum injection time was 86 ms, with HCD collision energy set to 65%. MS^3^ mass resolution (*R*) was set to 15,000 at *m/z* 400 for TMT6plex experiments, and 50,000 at *m/z* 400 for TMT11-plex experiments. Dynamic exclusion was set to exclude selected precursors for 60 s with a repeat count of 1. Nanospray voltage was set to 2.2 kV, with heated capillary temperature set to 300 °C, and an S-lens RF level equal to 30%. No sheath or auxiliary gas flow is applied.

#### Data Processing and Analysis

Acquired MS data was processed using Proteome Discoverer v. 2.2.0.388 software (Thermo) utilizing Mascot v 2.5.1 search engine (Matrix Science, London, UK) together with Percolator validation node for peptide-spectral match filtering ^47^. Data was searched against Uniprot protein database (canonical human and mouse sequences, EBI, Cambridge, UK) supplemented with sequences of common contaminants. Peptide search tolerances were set to 10 ppm for precursors, and 0.8 Da for fragments. Trypsin cleavage specificity (cleavage at K, R except if followed by P) allowed for up to 2 missed cleavages. Carbamidomethylation of cysteine was set as a fixed modification, methionine oxidation, and TMT-modification of N-termini and lysine residues were set as variable modifications. Data validation of peptide and protein identifications was done at the level of the complete dataset consisting of combined Mascot search results for all individual samples per experiment via the Percolator validation node in Proteome Discoverer. Reporter ion ratio calculations were performed using summed abundances with most confident centroid selected from 20 ppm window. Only peptide-to-spectrum matches that are unique assignments to a given identified protein within the total dataset are considered for protein quantitation. High confidence protein identifications were reported using a Percolator estimated <1% false discovery rate (FDR) cut-off. Differential abundance significance was estimated using a background-based ANOVA with Benjamini-Hochberg correction to determine adjusted p-values.

## Acknowledgements

We thank the members of the Nomura Research Group and Novartis Institutes for BioMedical Research for critical reading of the manuscript. This work was supported by Novartis Institutes for BioMedical Research and the Novartis-Berkeley Center for Proteomics and Chemistry Technologies (NB-CPACT) for all listed authors. This work was also supported by grants from the National Institutes of Health (F31CA225173 for CCW).

## References

(1) Burslem, G. M.; Crews, C. M. Small-Molecule Modulation of Protein Homeostasis. Chem. Rev. 2017, 117 (17), 11269–11301. https://doi.org/10.1021/acs.chemrev.7b00077.

(2) Lai, A. C.; Crews, C. M. Induced Protein Degradation: An Emerging Drug Discovery Paradigm. Nat. Rev. Drug Discov. 2017, 16 (2), 101–114. https://doi.org/10.1038/nrd.2016.211.

(3) Rape, M. Ubiquitylation at the Crossroads of Development and Disease. Nat. Rev. Mol. Cell Biol. 2018, 19 (1), 59–70. https://doi.org/10.1038/nrm.2017.83.

(4) Grossman, E. A.; Ward, C. C.; Spradlin, J. N.; Bateman, L. A.; Huffman, T. R.; Miyamoto, D. K.; Kleinman, J. I.; Nomura, D. K. Covalent Ligand Discovery against Druggable Hotspots Targeted by AntiCancer Natural Products. Cell Chem. Biol. 2017, 24 (11), 1368–1376.e4. https://doi.org/10.1016/j.chembiol.2017.08.013.

(5) Counihan, J. L.; Wiggenhorn, A. L.; Anderson, K. E.; Nomura, D. K. Chemoproteomics-Enabled Covalent Ligand Screening Reveals ALDH3A1 as a Lung Cancer Therapy Target. ACS Chem. Biol. 2018. https://doi.org/10.1021/acschembio.8b00381.

(6) Bateman, L. A.; Nguyen, T. B.; Roberts, A. M.; Miyamoto, D. K.; Ku, W.-M.; Huffman, T. R.; Petri, Y.; Heslin, M. J.; Contreras, C. M.; Skibola, C. F.; et al. Chemoproteomics-Enabled Covalent Ligand Screen Reveals a Cysteine Hotspot in Reticulon 4 That Impairs ER Morphology and Cancer Pathogenicity. Chem. Commun. Camb. Engl. 2017, 53 (53), 7234–7237. https://doi.org/10.1039/c7cc01480e.

(7) Backus, K. M.; Correia, B. E.; Lum, K. M.; Forli, S.; Horning, B. D.; González-Páez, G. E.; Chatterjee, S.; Lanning, B. R.; Teijaro, J. R.; Olson, A. J.; et al. Proteome-Wide Covalent Ligand Discovery in Native Biological Systems. Nature 2016, 534 (7608), 570–574. https://doi.org/10.1038/nature18002.

(8) Wang, C.; Weerapana, E.; Blewett, M. M.; Cravatt, B. F. A Chemoproteomic Platform to Quantitatively Map Targets of Lipid-Derived Electrophiles. Nat. Methods 2014, 11 (1), 79–85. https://doi.org/10.1038/nmeth.2759.

(9) Hacker, S. M.; Backus, K. M.; Lazear, M. R.; Forli, S.; Correia, B. E.; Cravatt, B. F. Global Profiling of Lysine Reactivity and Ligandability in the Human Proteome. Nat. Chem. 2017, 9 (12), 1181–1190. https://doi.org/10.1038/nchem.2826.

(10) Backus, K. M. Applications of Reactive Cysteine Profiling. Curr. Top. Microbiol. Immunol. 2018. https://doi.org/10.1007/82_2018_120.

(11) Liu, Y.; Patricelli, M. P.; Cravatt, B. F. Activity-Based Protein Profiling: The Serine Hydrolases. Proc. Natl. Acad. Sci. U. S. A. 1999, 96 (26), 14694–14699.

(12) Weerapana, E.; Wang, C.; Simon, G. M.; Richter, F.; Khare, S.; Dillon, M. B. D.; Bachovchin, D. A.; Mowen, K.; Baker, D.; Cravatt, B. F. Quantitative Reactivity Profiling Predicts Functional Cysteines in Proteomes. Nature 2010, 468 (7325), 790–795. https://doi.org/10.1038/nature09472.

(13) Roberts, A. M.; Miyamoto, D. K.; Huffman, T. R.; Bateman, L. A.; Ives, A. N.; Akopian, D.; Heslin, M. J.; Contreras, C. M.; Rape, M.; Skibola, C. F.; et al. Chemoproteomic Screening of Covalent Ligands Reveals UBA5 As a Novel Pancreatic Cancer Target. ACS Chem. Biol. 2017, 12 (4), 899–904. https://doi.org/10.1021/acschembio.7b00020.

(14) Wang, C.; Weerapana, E.; Blewett, M. M.; Cravatt, B. F. A Chemoproteomic Platform to Quantitatively Map Targets of Lipid-Derived Electrophiles. Nat. Methods 2014, 11 (1), 79–85. https://doi.org/10.1038/nmeth.2759.

(15) Anderson, K. E.; To, M.; Olzmann, J. A.; Nomura, D. K. Chemoproteomics-Enabled Covalent Ligand Screening Reveals a Thioredoxin-Caspase 3 Interaction Disruptor That Impairs Breast Cancer Pathogenicity. ACS Chem. Biol. 2017, 12 (10), 2522–2528. https://doi.org/10.1021/acschembio.7b00711.

(16) Vassilev, L. T.; Vu, B. T.; Graves, B.; Carvajal, D.; Podlaski, F.; Filipovic, Z.; Kong, N.; Kammlott, U.; Lukacs, C.; Klein, C.; et al. In Vivo Activation of the P53 Pathway by Small-Molecule Antagonists of MDM2. Science 2004, 303 (5659), 844–848. https://doi.org/10.1126/science.1092472.

(17) Schneekloth, A. R.; Pucheault, M.; Tae, H. S.; Crews, C. M. Targeted Intracellular Protein Degradation Induced by a Small Molecule: En Route to Chemical Proteomics. Bioorg. Med. Chem. Lett. 2008, 18 (22), 5904–5908. https://doi.org/10.1016/j.bmcl.2008.07.114.

(18) Malecka, K. A.; Fera, D.; Schultz, D. C.; Hodawadekar, S.; Reichman, M.; Donover, P. S.; Murphy, M. E.; Marmorstein, R. Identification and Characterization of Small Molecule Human Papillomavirus E6 Inhibitors. ACS Chem. Biol. 2014, 9 (7), 1603–1612. https://doi.org/10.1021/cb500229d.

(19) Staudinger, J. L. The Molecular Interface Between the SUMO and Ubiquitin Systems. Adv. Exp. Med. Biol. 2017, 963, 99–110. https://doi.org/10.1007/978-3-319-50044-7_6.

(20) Sriramachandran, A. M.; Dohmen, R. J. SUMO-Targeted Ubiquitin Ligases. Biochim. Biophys. Acta 2014, 1843 (1), 75–85. https://doi.org/10.1016/j.bbamcr.2013.08.022.

(21) Tan, B.; Mu, R.; Chang, Y.; Wang, Y.-B.; Wu, M.; Tu, H.-Q.; Zhang, Y.-C.; Guo, S.-S.; Qin, X.-H.; Li, T.; et al. RNF4 Negatively Regulates NF-?B Signaling by down-Regulating TAB2. FEBS Lett. 2015, 589 (19 Pt B), 2850–2858. https://doi.org/10.1016/j.febslet.2015.07.051.

(22) Fryrear, K. A.; Guo, X.; Kerscher, O.; Semmes, O. J. The Sumo-Targeted Ubiquitin Ligase RNF4 Regulates the Localization and Function of the HTLV-1 Oncoprotein Tax. Blood 2012, 119 (5), 1173–1181. https://doi.org/10.1182/blood-2011-06-358564.

(23) Bilodeau, S.; Caron, V.; Gagnon, J.; Kuftedjian, A.; Tremblay, A. A CK2-RNF4 Interplay Coordinates Non-Canonical SUMOylation and Degradation of Nuclear Receptor FXR. J. Mol. Cell Biol. 2017, 9 (3), 195–208. https://doi.org/10.1093/jmcb/mjx009.

(24) Filippakopoulos, P.; Qi, J.; Picaud, S.; Shen, Y.; Smith, W. B.; Fedorov, O.; Morse, E. M.; Keates, T.; Hickman, T. T.; Felletar, I.; et al. Selective Inhibition of BET Bromodomains. Nature 2010, 468 (7327), 1067–1073. https://doi.org/10.1038/nature09504.

(25) Zengerle, M.; Chan, K.-H.; Ciulli, A. Selective Small Molecule Induced Degradation of the BET Bromodomain Protein BRD4. ACS Chem. Biol. 2015, 10 (8), 1770–1777. https://doi.org/10.1021/acschembio.5b00216.

(26) Winter, G. E.; Buckley, D. L.; Paulk, J.; Roberts, J. M.; Souza, A.; Dhe-Paganon, S.; Bradner, J. E. DRUG DEVELOPMENT. Phthalimide Conjugation as a Strategy for in Vivo Target Protein Degradation. Science 2015, 348 (6241), 1376–1381. https://doi.org/10.1126/science.aab1433.

(27) Gadd, M. S.; Testa, A.; Lucas, X.; Chan, K.-H.; Chen, W.; Lamont, D. J.; Zengerle, M.; Ciulli, A. Structural Basis of PROTAC Cooperative Recognition for Selective Protein Degradation. Nat. Chem. Biol. 2017, 13 (5), 514–521. https://doi.org/10.1038/nchembio.2329.

(28) Nowak, R. P.; DeAngelo, S. L.; Buckley, D.; He, Z.; Donovan, K. A.; An, J.; Safaee, N.; Jedrychowski, M. P.; Ponthier, C. M.; Ishoey, M.; et al. Plasticity in Binding Confers Selectivity in Ligand-Induced Protein Degradation. Nat. Chem. Biol. 2018. https://doi.org/10.1038/s41589-018-0055-y.

(29) Weerapana, E.; Wang, C.; Simon, G. M.; Richter, F.; Khare, S.; Dillon, M. B. D.; Bachovchin, D. A.; Mowen, K.; Baker, D.; Cravatt, B. F. Quantitative Reactivity Profiling Predicts Functional Cysteines in Proteomes. Nature 2010, 468 (7325), 790–795. https://doi.org/10.1038/nature09472.

(30) Kokosza, K.; Balzarini, J.; Piotrowska, D. G. Novel 5-Arylcarbamoyl-2-Methylisoxazolidin-3-Yl-3-Phosphonates as Nucleotide Analogues. Nucleosides Nucleotides Nucleic Acids 2014, 33 (8), 552–582. https://doi.org/10.1080/15257770.2014.909046.

(31) Talaty, E. R.; Young, S. M.; Dain, R. P.; Stipdonk, M. J. V. A Study of Fragmentation of Protonated Amides of Some Acylated Amino Acids by Tandem Mass Spectrometry: Observation of an Unusual Nitrilium Ion. Rapid Commun. Mass Spectrom. 2011, 25 (9), 1119–1129. https://doi.org/10.1002/rcm.4965.

(32) Timokhin, V. I.; Gastaldi, S.; Bertrand, M. P.; Chatgilialoglu, C. Rate Constants for the β-Elimination of Tosyl Radical from a Variety of Substituted Carbon-Centered Radicals. J. Org. Chem. 2003, 68 (9), 3532–3537. https://doi.org/10.1021/jo026870b.

(33) Cee, V. J.; Volak, L. P.; Chen, Y.; Bartberger, M. D.; Tegley, C.; Arvedson, T.; McCarter, J.; Tasker, A. S.; Fotsch, C. Systematic Study of the Glutathione (GSH) Reactivity of N-Arylacrylamides: 1. Effects of Aryl Substitution. J. Med. Chem. 2015, 58 (23), 9171–9178. https://doi.org/10.1021/acs.jmedchem.5b01018.

(34) Le Sann, C.; Huddleston, J.; Mann, J. Synthesis and Preliminary Evaluation of Novel Analogues of Quindolines as Potential Stabilisers of Telomeric G-Quadruplex DNA. Tetrahedron 2007, 63 (52), 12903–12911. https://doi.org/10.1016/j.tet.2007.10.045.

(35) Ikoma, M.; Oikawa, M.; Sasaki, M. Synthesis and Domino Metathesis of Functionalized 7-Oxanorbornene Analogs toward Cis-Fused Heterocycles. Tetrahedron 2008, 64 (12), 2740–2749. https://doi.org/10.1016/j.tet.2008.01.067.

(36) Cho, S.-D.; Park, Y.-D.; Kim, J.-J.; Lee, S.-G.; Ma, C.; Song, S.-Y.; Joo, W.-H.; Falck, J. R.; Shiro, M.; Shin, D.-S.; et al. A One-Pot Synthesis of Pyrido[2,3-b][1,4]Oxazin-2-Ones. J. Org. Chem. 2003, 68 (20), 7918–7920. https://doi.org/10.1021/jo034593i.

(37) Magolan, J.; Carson, C. A.; Kerr, M. A. Total Synthesis of (±)-Mersicarpine. Org. Lett. 2008, 10 (7), 1437–1440. https://doi.org/10.1021/ol800259s.

(38) Louie, S. M.; Grossman, E. A.; Crawford, L. A.; Ding, L.; Camarda, R.; Huffman, T. R.; Miyamoto, D. K.; Goga, A.; Weerapana, E.; Nomura, D. K. GSTP1 Is a Driver of Triple-Negative Breast Cancer Cell Metabolism and Pathogenicity. Cell Chem. Biol. 2016, 23 (5), 567–578. https://doi.org/10.1016/j.chembiol.2016.03.017.

(39) Medina-Cleghorn, D.; Bateman, L. A.; Ford, B.; Heslin, A.; Fisher, K. J.; Dalvie, E. D.; Nomura, D. K. Mapping Proteome-Wide Targets of Environmental Chemicals Using Reactivity-Based Chemoproteomic Platforms. Chem. Biol. 2015, 22 (10), 1394–1405. https://doi.org/10.1016/j.chembiol.2015.09.008.

(40) Groocock, L. M.; Nie, M.; Prudden, J.; Moiani, D.; Wang, T.; Cheltsov, A.; Rambo, R. P.; Arvai, A. S.; Hitomi, C.; Tainer, J. A.; et al. RNF4 Interacts with Both SUMO and Nucleosomes to Promote the DNA Damage Response. EMBO Rep. 2014, 15 (5), 601–608. https://doi.org/10.1002/embr.201338369.

(41) Schrödinger Release 2018-1: Maestro, Schrödinger, LLC, New York, NY, 2018.

(42) Zhu, K.; Borrelli, K. W.; Greenwood, J. R.; Day, T.; Abel, R.; Farid, R. S.; Harder, E. Docking Covalent Inhibitors: A Parameter Free Approach To Pose Prediction and Scoring. J. Chem. Inf. Model. 2014, 54 (7), 1932–1940. https://doi.org/10.1021/ci500118s.

(43) Jessani, N.; Humphrey, M.; McDonald, W. H.; Niessen, S.; Masuda, K.; Gangadharan, B.; Yates, J. R.; Mueller, B. M.; Cravatt, B. F. Carcinoma and Stromal Enzyme Activity Profiles Associated with Breast Tumor Growth in Vivo. Proc. Natl. Acad. Sci. U. S. A. 2004, 101 (38), 13756–13761. https://doi.org/10.1073/pnas.0404727101.

(44) Smith, P. K.; Krohn, R. I.; Hermanson, G. T.; Mallia, A. K.; Gartner, F. H.; Provenzano, M. D.; Fujimoto, E. K.; Goeke, N. M.; Olson, B. J.; Klenk, D. C. Measurement of Protein Using Bicinchoninic Acid. Anal. Biochem. 1985, 150 (1), 76–85.

(45) Xu, T.; Park, S. K.; Venable, J. D.; Wohlschlegel, J. A.; Diedrich, J. K.; Cociorva, D.; Lu, B.; Liao, L.; Hewel, J.; Han, X.; et al. ProLuCID: An Improved SEQUEST-like Algorithm with Enhanced Sensitivity and Specificity. J. Proteomics 2015, 129, 16–24. https://doi.org/10.1016/j.jprot.2015.07.001.

(46) Thomas, J. R.; Brittain, S. M.; Lipps, J.; Llamas, L.; Jain, R. K.; Schirle, M. A Photoaffinity Labeling-Based Chemoproteomics Strategy for Unbiased Target Deconvolution of Small Molecule Drug Candidates. Methods Mol. Biol. Clifton NJ 2017, 1647, 1–18. https://doi.org/10.1007/978-1-4939-7201-2_1.

(47) Käll, L.; Canterbury, J. D.; Weston, J.; Noble, W. S.; MacCoss, M. J. Semi-Supervised Learning for Peptide Identification from Shotgun Proteomics Datasets. Nat. Methods 2007, 4 (11), 923–925. https://doi.org/10.1038/nmeth1113.

